# Predicting compatibility between ferredoxins and the Fe protein of nitrogenase using *in silico* protein modeling

**DOI:** 10.1101/2025.08.21.671581

**Authors:** Adity Biswas, Katerina Trachtova, Kathryn R. Fixen

## Abstract

Biological nitrogen fixation is the process by which certain bacteria and archaea use the enzyme nitrogenase to reduce atmospheric nitrogen into bioavailable ammonium. Engineering non-nitrogen-fixing organisms, like plants, to use nitrogenase could reduce dependency on synthetic fertilizer and mitigate the environmental impacts of industrial fertilizer production. However, nitrogenase activity requires delivery of reducing power by small electron carrying proteins known as ferredoxins and flavodoxins, and successfully engineering nitrogenase into new systems will require a mechanistic understanding of electron delivery by these proteins. Most organisms often have multiple ferredoxins, raising the question of which ferredoxin can support nitrogenase activity. The purpose of this study is to gain insight into how we can predict which ferredoxin is compatible with the Fe protein, the component of nitrogenase that interacts with ferredoxin or flavodoxin. Our *in silico* protein-protein docking simulations reveal that ferredoxins and flavodoxins involved in nitrogen fixation have a distance ≤ 10 Å between their redox cofactor and the [4Fe-4S] cluster of the Fe protein. We found shorter cofactor distance contributes to faster intermolecular electron tunneling rates (> 10^6^ sec^-1^). These nitrogen-fixing bacterial ferredoxins also form more complementary interactions with the Fe protein compared to non-nitrogen-fixing bacterial and plant ferredoxins. Heterologous expression of a set of ferredoxins from both nitrogen-fixing and non-nitrogen-fixing bacteria in the diazotroph *Rhodopseudomonas palustris* support our model-derived prediction that cofactor distances of ≤ 10 Å favor nitrogenase compatibility. These findings offer a framework to predict and potentially enhance ferredoxin–nitrogenase compatibility, which will help to improve our ability to engineer nitrogen fixation into non-nitrogen-fixing organisms like plants.

## Introduction

Biological nitrogen fixation describes the reduction of atmospheric nitrogen gas (N2) into bioavailable ammonium (NH4) by the oxygen-sensitive enzyme nitrogenase. Biological nitrogen fixation is a restricted metabolic trait found in specific bacterial and archaeal species and was recently found to occur in a marine alga, *Braarudosphaera bigelowii* [1–4]. Historically, limited fixed nitrogen availability constrained crop production until the development of the industrial Haber-Bosch process, which produces 200 million tons of fertilizers annually to support global agriculture [5, 6]. However, this process has significant environmental costs, including greenhouse gas emissions and eutrophication from fertilizer runoff [7, 8]. One alternative to reduce reliance on synthetic fertilizer involves engineering plants to fix nitrogen using nitrogenase [9]. Recent advancements in identifying the minimal gene set required for nitrogen fixation and successful expression of nitrogenase components in eukaryotic organelles, such as chloroplasts and mitochondria, highlight significant progress in building a functional nitrogenase in non-nitrogen-fixing organisms [7, 10–16].

Nitrogenase is composed of two subunits, a homodimeric NifH subunit known as the iron (Fe) protein and a heterotetrameric NifDK subunit known as the iron-molybdenum (MoFe) protein [17]. Over eight electron-transferring catalytic cycles, the Fe protein hydrolyzes 16 ATP and delivers 8 electrons to the MoFe protein to reduce one molecule of N2 to two moles of ammonia and one mole of hydrogen. The Fe protein uses electrons donated by low potential protein electron carriers such as ferredoxin or flavodoxin [18, 19]. Both the Fe protein and the MoFe protein have been successfully expressed in plant and yeast organelles, suggesting that active nitrogenase could potentially be assembled in these compartments [15, 20–24]. However, successful integration of nitrogenase into non-nitrogen-fixing organisms, including plants, will also require understanding of the properties of ferredoxins to deliver electrons to the Fe protein. Previous studies have demonstrated that plant ferredoxins, including those from the plant organelles chloroplast, mitochondria, and root plastid, can support some nitrogenase activity in *Escherichia coli* engineered to express nitrogenase [25]. However, the molecular underpinnings that led to the variability in nitrogenase activity supported by these ferredoxins remains unclear.

Studies on electron transfer between electron-carrying cofactors within an enzyme complex have shown that there are two properties that natural selection has acted on to control electron tunneling between the cofactors: redox potential and the distance between the cofactors [26–32]. The reported redox potential of the ADP-bound Fe protein varies from -415 mV [33] to -470 mV [18]. Based on the midpoint potential of the Fe protein, we would expect nitrogen-fixing ferredoxins and flavodoxins to have redox potentials lower than the Fe protein. However, some nitrogen-fixing ferredoxins and flavodoxins such as FdxH from *Anabaena* (*Nostoc*) *sp.* PCC7120 (*As*FdxH) (-351 mV [34]) and NifF of *Klebsiella oxytoca* (*Ko*NifF) (-412 mV [35]) have redox potentials higher than the Fe protein while some ferredoxins such as Fer1 from *R. palustris* (*Rp*Fer1) (- 452, -583 mV [36]) have very low redox potentials. Additionally, all plant ferredoxins tested in the *E. coli* heterologous nitrogen-fixing system could support some nitrogenase activity, yet there was no correlation between the reported redox potential for the ferredoxin with the magnitude of nitrogenase activity observed [25]. This suggests that redox potential alone may not be sufficient to predict which ferredoxins are compatible with the Fe protein.

Although the Fe protein and its cognate electron carriers interact transiently, we hypothesized that co-evolution has likely positioned their redox cofactors close enough to enable efficient electron transfer. To test this hypothesis, we simulated protein-protein docking models of the Fe protein from our chassis diazotroph *Rhodopseudomonas palustris* complexed with electron carriers from both nitrogen-fixing bacteria (**Supplemental table 1**) and non-nitrogen-fixing bacteria (**Supplemental table 2**) and plants (**Supplemental table 3**) [37]. We measured the edge-to-edge distance between redox cofactors of the partner proteins, and we found that cofactor distance between non-nitrogen-fixing ferredoxins is larger (> 10 Å) compared to nitrogen-fixing ferredoxins and flavodoxins. The increased distance between the iron-sulfur (FeS) clusters results in slower electron tunneling rates, even when the redox potential of the ferredoxin is lower than -400 mV. We show the distance between the electron-carrying cofactors is a better indicator of compatibility between a ferredoxin and the Fe protein than redox potential alone. These findings suggest that shorter distances between FeS clusters of a ferredoxin and the Fe protein could enhance the ability of a plant ferredoxin to support nitrogenase activity.

## Results

### Nitrogen-fixing electron carriers have the shortest distance between their redox cofactor and the FeS cluster of the Fe protein

We modeled the distance between electron-carrying cofactors in the ferredoxin/flavodoxin and the Fe protein, as distance is known to influence electron transfer and may serve as a predictive parameter for assessing ferredoxin and flavodoxin compatibility with the Fe protein [29]. The distance between electron-carrying cofactors may also reflect how well a ferredoxin or flavodoxin interacts with the Fe protein since this interface has likely been optimized over evolutionary time to bring the electron-carrying cofactors closer together. We constructed *in silico* protein-protein docking models using the Fe protein from *R. palustris* paired with either a ferredoxin or flavodoxin from nitrogen-fixing bacteria (**Supplemental 1**), non-nitrogen-fixing bacteria (**Supplemental table 2**), and plants (**Supplemental table 3**). The ADP-bound Fe protein of *R. palustris* was simulated using a template-directed model (PDB: 1FP6) of the ADP-bound Fe protein of *Azotobacter vinelandii* [38]. We used ClusPro2.0 to create docking models between electron carriers and the ADP-bound Fe protein [37]. We then analyzed these models to assess the role of distance between the electron carrying cofactor of ferredoxin or flavodoxin and the [4Fe-4S] cluster of the Fe protein. We measured the edge-to-edge distance, *R* (Å), between the [4Fe-4S] cluster of ADP-bound Fe protein and either the FeS cluster in a ferredoxin or the FMN in a flavodoxin (**Fig. 1**).

**Fig. 1.**
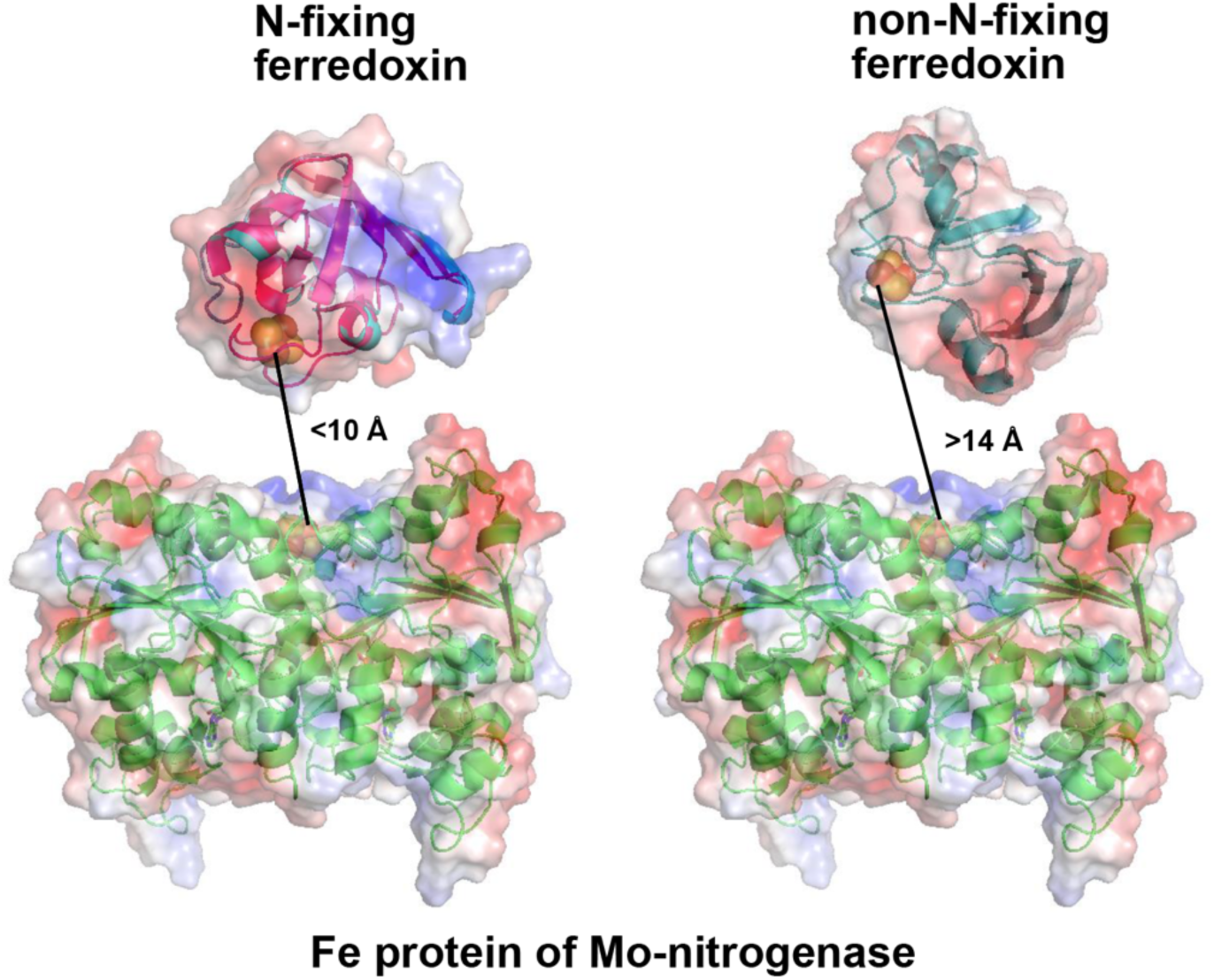
*In silico* protein modeling between a ferredoxin and the Fe protein of nitrogenase. A nitrogen-fixing bacterial ferredoxin like FdxH from *Anabaena* (*Nostoc*) *sp.* PCC 7120 (UniProt: P11053) forms complementary docking interactions with a model of the Fe protein (WP_011160152) of nitrogenase from *Rhodopseudomonas palustris* that brings the FeS cofactors within 10 Å. While non-nitrogen fixing plant ferredoxins such as FD3 from *Arabidopsis thaliana* (NP_180320.1) show edge-to-edge cofactor distances greater than 14 Å.

Broadly, the average cofactor distances between the nitrogen-fixing electron carriers *Rp*Fer1, *Rp*FldA, *Rp*FerN, *Ko*NifF, *Rc*FdN, *Rc*FdA, and *As*FdxH and the Fe protein had the shortest distances, and distances were shorter than 10 Å, which is consistent with the 6.4 Å previously observed between the flavodoxin NifF and the ADP-bound Fe protein of *Azotobacter vinelandii* [39] (**Fig. 2 and Supplemental Table 1**). *In vivo* studies confirmed that the absence of these nitrogen-fixing ferredoxins and flavodoxins showed a strong growth defect in their native organisms in nitrogen-fixing conditions [40–42]. This evidence supports shorter average cofactor distance observed for nitrogen-fixing ferredoxins reflects that they have evolved over time to interact more efficiently and specifically with the Fe protein.

**Fig. 2.**
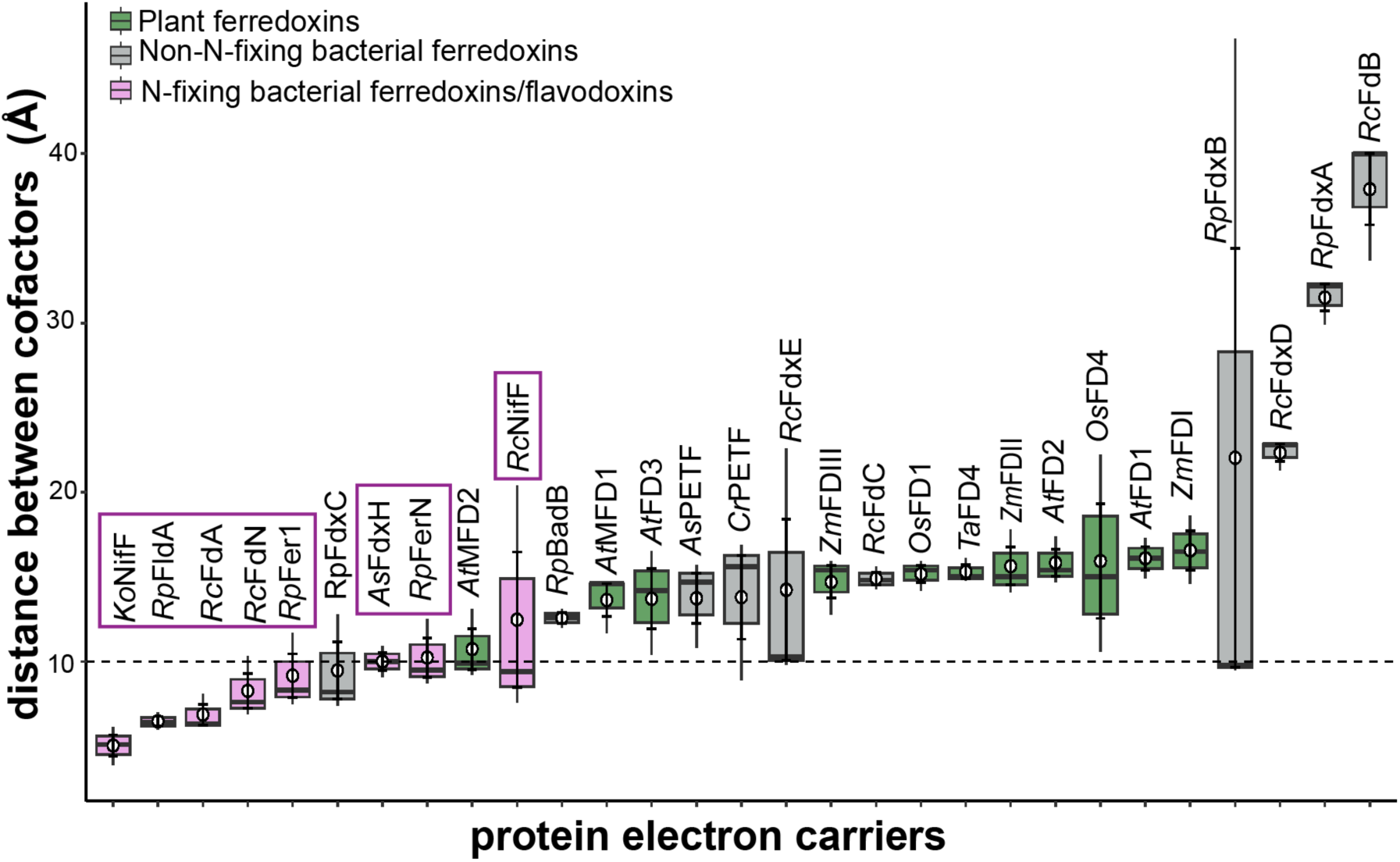
Nitrogen fixing electron carriers like ferredoxins and flavodoxins have a shorter cofactor distance from the Fe protein of nitrogenase in docking simulations. Edge-to-edge cofactor distance measured in simulated interactions between nitrogen-fixing bacterial ferredoxins/flavodoxins (pink), non-nitrogen-fixing bacterial ferredoxins (gray), plant ferredoxins (green), and the modelled nitrogenase Fe protein from *R. palustris*. Electron carriers involved in nitrogen fixation include *R. palustris* (*Rp*Fer1, *Rp*FldA, *Rp*FerN), *R. capsulatus* (*Rc*FdN, *Rc*NifF, *Rc*FdA), *K. oxytoca* (*Ko*nifF), and *Anabaena sp.* PCC 7120 (*As*FdxH) highlighted with pink box. The non-N fixing ferredoxins from bacteria include *R. palustris* (*Rp*BadB, *Rp*FdxA, *Rp*FdxB), *R. capsulatus* (*Rc*FdxB, *Rc*FdxC, *Rc*FdxD), *Anabaena* (*Nostoc*) *sp.* PCC 7120 (*As*PETF), and *C. reinhardtii* (*Cr*PETF). Plant ferredoxins include chloroplast ferredoxins from *A. thaliana* (*At*FD1 and *At*FD2), *Z. mays* (*Zm*FDI and *Zm*FDII), *O. sativa* (*Os*FD1), *T. aestivum* (*Ta*FD4); root plastid ferredoxins from *A. thaliana* (*At*FD3), *Z. mays* (*Zm*FDIII), *O. sativa* (*Os*FD4); and mitochondrial ferredoxins from *A. thaliana* (*At*MFD1, *At*MFD2). Dashed line at 10 Å represents the threshold under which most nitrogen-fixing electron carriers fall. Data are from three independent docking simulations. For each electron carrier, the central line in the box shows the median cofactor distance measured, and the edges of the box represent the 25^th^ and 75^th^ percentiles and whiskers extend to the minimum and maximum cofactor distances within 1.5 interquartile ranges. Distance data for each model is listed in **Supplemental Table 1 to 3**.

In contrast, the distances between cofactors were much larger for non-nitrogen-fixing ferredoxins both from bacteria and plants compared to ferredoxins known to be involved in nitrogen fixation (**Fig. 2**). The average cofactor distance for non-nitrogen-fixing bacterial ferredoxins was higher than 14 Å except for FdxC from *R. palustris* (*Rp*FdxC) that had an average cofactor distance of 9.5 Å (**Fig. 2 and Supplemental Table 2**). *Rp*FdxC has not been shown to play a role in nitrogen fixation in *R. palustris,* but its docking model proposed that it had less than 10 Å between its [4Fe-4S] cluster and the [4Fe-4S] cluster of the Fe protein, a distance similar to other ferredoxins involved in nitrogen fixation (**Fig. 2 and Supplemental Table 1**). We found *Rp*FdxC has 74% amino acid identity to *Rc*FdA, a ferredoxin from *R. capsulatus*, which is not the main electron donor to nitrogenase in nitrogen-fixing conditions but acts as an electron donor for nitrogenase *in vitro* [43]. However, *Rp*FdxC and *Rc*FdA are required for growth under non-nitrogen-fixing conditions, suggesting they serve additional roles [44, 45]. These results demonstrate that protein modeling can provide insight into potential functional interactions between electron carriers and nitrogenase.

Similarly, the average cofactor distances for plant ferredoxins found in the chloroplast, root plastid, or mitochondria were also greater than 14 Å (**Fig. 2 and Supplemental Table 3**). The exception being the mitochondrial ferredoxin, *At*MFD2, which had an average cofactor distance of 10.7 Å (**Fig. 2 and Supplemental Table 1**). We analyzed relative nitrogenase activity for plant ferredoxins reported in an engineered *E. coli* with our measured average cofactor distances. This time, we found a significant correlation (R^2^ = 0.644) between cofactor distance and nitrogenase activity reported in [25]. These results suggest plant ferredoxins supported less nitrogenase activity because they had larger cofactor distance. Our findings indicate shorter cofactor distance (≤ 10 Å) between a ferredoxin and the Fe protein is important to support nitrogenase activity.

### Nitrogen-fixing ferredoxins have the fastest calculated electron tunneling rate

While the distance between the electron-carrying cofactors can indicate how well two proteins interact, the rate of electron tunneling between them depends on both the difference between the redox potentials and the distance between redox cofactors [46]. Electron tunneling rate was calculated for each electron carrier protein and the Fe protein to determine how differences in distance and redox potential impacted electron tunneling between the Fe protein and its electron donor. Two empirical formulas (**Eq. 1 and 2 in Materials and Methods**) were used to calculate the electron tunneling rate based on either exergonic or endergonic driving force [30]. Ferredoxins and flavodoxins previously shown to be involved in nitrogen fixation had the fastest electron tunneling rates, >10^6^ sec^-1^ (**Fig. 3A and Supplemental Table 1**). This was true even for NifF from *Klebsiella* and FdxH from *Anabaena*, which have a higher midpoint potential than the Fe protein [34, 35]. Although electron carriers from *Klebsiella* and *Anabaena* share a similar midpoint potential as some of the plant ferredoxins being investigated, their electron tunneling rates are much faster because their cofactors come within 10 Å of the [4Fe-4S] cluster of the Fe protein. In a natural electron tunneling path, shorter distances between cofactors support faster electron tunneling rates, consistent with our observations for nitrogen-fixing ferredoxins and flavodoxins, even under thermodynamically unfavorable conditions [28].

**Fig. 3.**
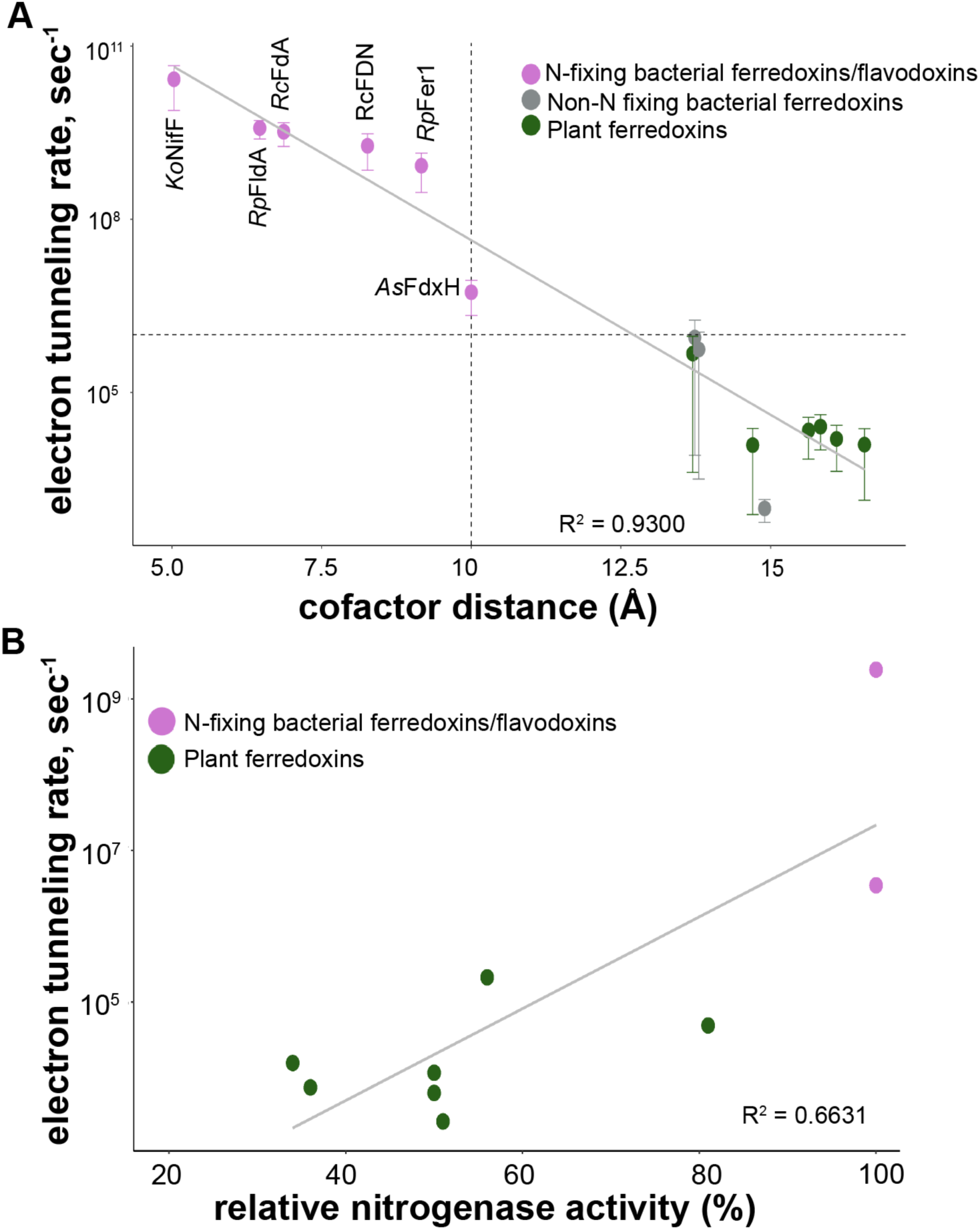
Nitrogen-fixing ferredoxins and flavodoxins have faster calculated electron tunneling rates than non-nitrogen-fixing electron carriers. (A) The mean electron tunneling rate increases as distance between the electron-carrying cofactors decreases. Ferredoxins and flavodoxins involved in nitrogen fixation (pink circles) had the fastest electron tunneling rates compared to other non-nitrogen-fixing ferredoxins from bacteria (grey circles) and plants (black circles). Error bars represent one standard deviation from the mean. Dashed line on X-axis represents a cofactor distance of 10 Å while the dashed line on Y-axis represents an electron tunneling rate of 10^6^ sec^-1^. (B) There is a positive correlation between the calculated electron tunneling rate and the relative nitrogenase activity reported in [25]. Ferredoxin and flavodoxin involved in nitrogen fixation are represented by pink circles and plant ferredoxins are shown in black. Calculated electron tunneling rates used in (A) and (B) can also be found in **Supplemental Table 1-3**.

In contrast, we found non-nitrogen-fixing bacterial and plant ferredoxins had much slower electron tunneling rates, ≤ 10^6^ sec^-1^, with an average of 10^4^ sec^-1^ (**Fig. 3A and Supplemental Table 2 and 3**). Among plant ferredoxins, the root ferredoxin *At*FD3 has the fastest electron tunneling rate, 10^5^ sec^-1^. Similar to electron carriers from *Klebsiella* and *Anabaena* described above, *At*Fd3 has a higher redox potential (-337 mV) but has one of the shortest modeled cofactor distance among non-nitrogen-fixing ferredoxins (13.7 Å) [48]. We analyzed relative nitrogenase activity reported [25] with our calculated electron tunneling rates for each plant ferredoxins to determine if a slower electron tunneling rate could impact *in vivo* nitrogenase activity (**Fig. 3B**). We found there is a correlation (R^2^=0.663) between the calculated electron tunneling rate and the nitrogenase activity reported (**Fig. 3B**). This suggests that ferredoxins that have a slower electron tunneling rate support less *in vivo* nitrogenase activity.

To investigate how a plant ferredoxin could be engineered to enhance electron transfer to nitrogenase, we modeled electron tunneling rates across a range of redox potentials (from –1000 mV to –200 mV, in 25 mV increments) and cofactor distances (from 4 Å to 20 Å, in 1 Å increments) (**Fig. 4**). We assumed that a tunneling rate of at least 10^6^ sec^-1^ was necessary to maintain nitrogenase activity since this was the slowest rate calculated for a nitrogen-fixing ferredoxin (**Fig. 3A**). To achieve this rate, the cofactor distance between the FeS clusters of the Fe protein and a plant ferredoxin like *Zm*Fd3, which has a redox potential of -321 mV, would need to decrease from 14.7 Å to 10 Å. However, if we assume that the distance between the *Zm*Fd3 and the Fe protein stays the same, the redox potential of the *Zm*Fd3 would have to drop to -850 mV to achieve a similar tunneling rate at 10^6^ sec^-1^. Since no known [2Fe-2S] cluster ferredoxin has a redox potential that low, engineering a non-nitrogen-fixing ferredoxin with a shorter cofactor distance may be a more practical strategy to increase the electron tunneling rate from a plant ferredoxin to nitrogenase.

**Fig. 4.**
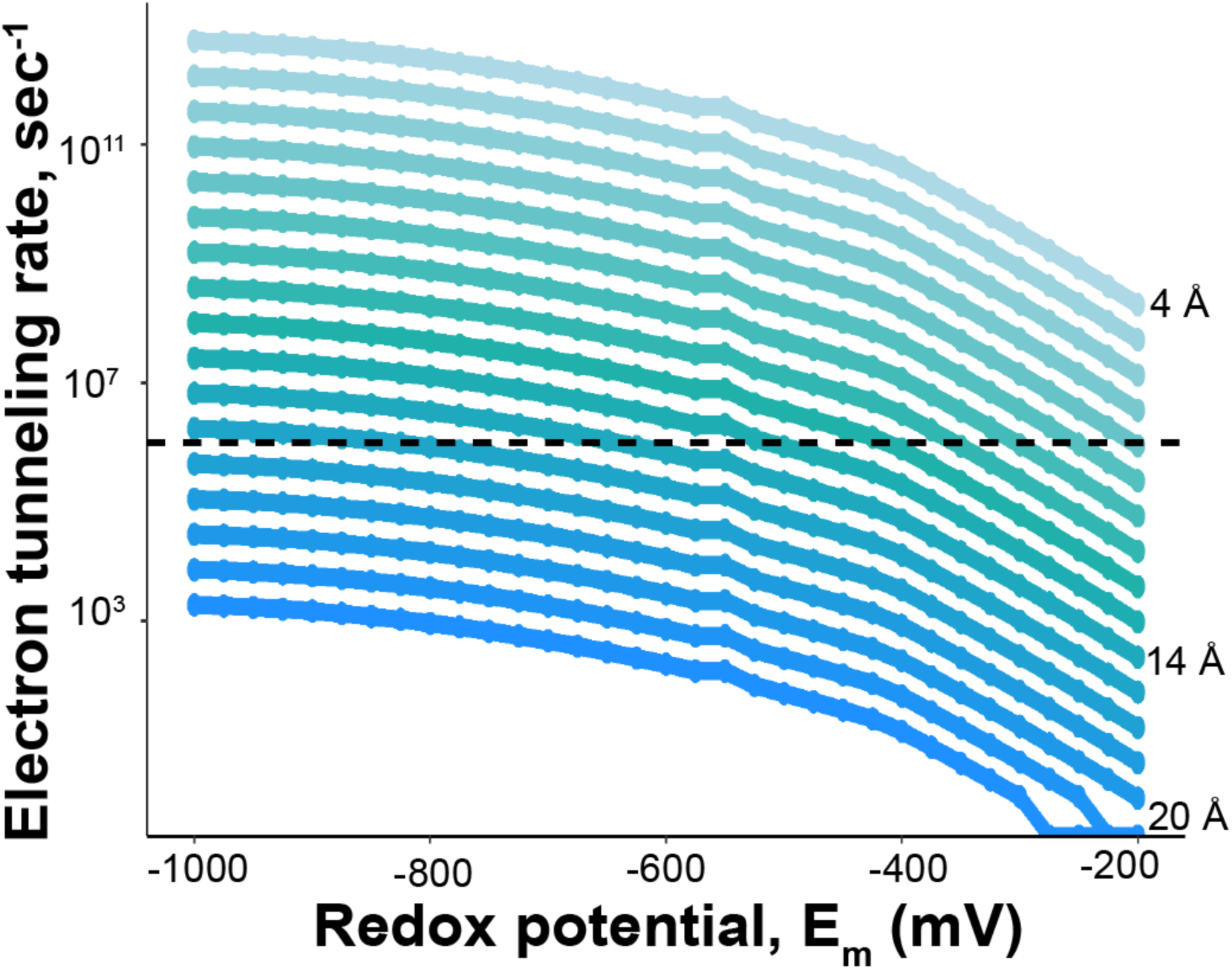
Modeling of calculated electron tunneling rates as a function of cofactor distance and redox potential. The electron tunneling rate at a given potential from - 1000 mV to -200 mV was determined with increasing distance between the cofactors. The shortest distance was 4 Å (top, light blue), and each line represents an increase in distance of 1 Å up to 20 Å (bottom, dark blue line). The dashed line reflects an electron tunneling rate 10^6^ sec^-1^.

### Electrostatic interactions at the protein-protein interface may bring the FeS clusters in a ferredoxin and the Fe protein closer together

Our *in silico* protein modeling suggests a cofactor distance ≤ 10 Å is a critical determinant for supporting nitrogenase activity. Therefore, evaluating the docking interface between electron carriers and the Fe protein is essential to identify the factors that influence cofactor distance. Electrostatic interactions at the interface of NifF and the Fe protein in *A. vinelandii* are known to be important for their interaction [39]. Thus, we analyzed the apparent electrostatic interactions between each modelled electron carrier and the model of the *R. palustris* Fe protein. Charged residues including arginine 100, arginine 140, glutamate 59, and glutamate 104 on the Fe protein were identified in chemical crosslinking and *in silico* docking experiments when interacting with NifF of *A. vinelandii* [39]. Our docking models with nitrogen-fixing ferredoxins/flavodoxins suggest the corresponding charged residues arginine 101 (R101), arginine 140 (R140), glutamate 112 (E112), and glutamate 69 (E69) on the Fe protein of *R. palustris* are also important to form salt bridge interactions (**Fig. 5**). We performed multiple sequence alignment of homologous Fe proteins from selected diazotrophs from clade I (Nif I), which includes Mo-nitrogenase from facultative anaerobes; clade II (Nif II), which includes Mo-nitrogenase from obligate anaerobes; and clade III (Nif III), which includes alternative nitrogenases from both facultative and obligate anaerobes, of the nitrogenase phylogenetic tree [48] (**Supplemental Fig. 1**). Alignment of these homologous Fe protein sequences suggests R101, R140, E112 are 100% conserved and E69 is 70% conserved or is replaced with another negatively charged amino acid like an aspartate (D) (**Supplemental Fig. 1**). This result suggests charged residues and salt-bridge interactions are a conserved feature of interactions between nitrogen-fixing electron carriers and the Fe protein.

**Fig. 5.**
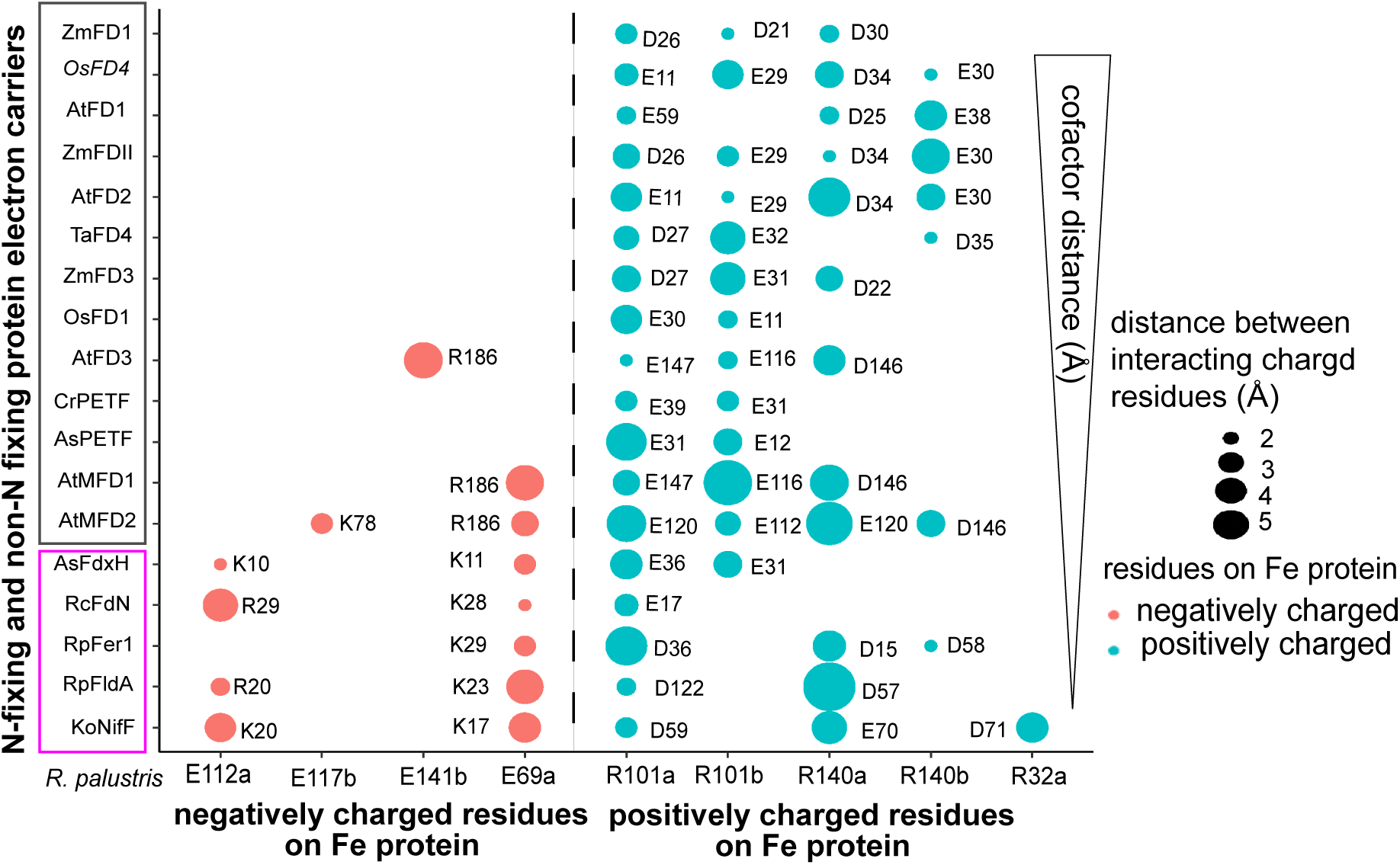
Electron carriers involved in nitrogen fixation have more electrostatic interactions with the Fe protein of nitrogenase. Electrostatic interactions at the docking interface between electron carriers and Fe protein of nitrogenase arranged by the distance between the [4Fe-4S] cluster of the Fe protein and the electron transferring cofactor of the ferredoxin or flavodoxin. The interacting charged residues from each electron carrier protein is shown next to a bubble representing the distance between the residue on the protein electron carrier and the corresponding residue on the Fe protein from *R. palustris* shown on the x-axis. Residues in the protein electron carrier that interact with negatively charged residues of the Fe protein are shown in red and interactions with positive residues of the Fe protein are shown in blue. Because the Fe protein is a homodimer, a and b are used to indicate whether a bond is being made to the same amino acid on the arbitrarily selected chain a or b of the Fe protein dimer.

To investigate how surface charge of the ferredoxin plays a role in forming salt-bridge interactions, we compared the individual protein models of nitrogen-fixing ferredoxins and non-nitrogen-fixing ferredoxins. From individual protein models of nitrogen-fixing ferredoxins, we found they generally have a cluster of positively charged amino acids and a cluster of negatively charged amino acids on their surface. A similar surface charge pattern on the protein structure of nitrogen-fixing protein electron carriers has been observed by others [39, 42, 49]. We found the surface charge distribution on nitrogen-fixing ferredoxins facilitates the formation of salt-bridge interactions with the Fe protein (**Fig. 5**). This suggest that the differences in surface charge observed for nitrogen-fixing ferredoxins and flavodoxins enables more salt-bridge interactions, bringing the cofactors closer together.

To further understand the role of charged amino acids, we substituted lysine 10 and 11 with glutamate (K10E) and alanine (K11A) in FdxH from *Anabaena* and did docking simulations with the Fe protein of *R. palustris*. We found the alteration of these residues increased the average cofactor distance from 10 Å to 15.3 Å in our docking models. Use of this variant of FdxH (*As*FdxH^K10E,K11A^) instead of wild-type FdxH *in vitro* resulted in a 31% reduction in nitrogenase activity [42]. This result suggests charge compatibility influences the distance between FeS clusters, and hence, nitrogenase activity. Plant ferredoxins have mostly negatively charged amino acids on their protein surface and form fewer salt-bridge interactions with the Fe protein (**Fig. 5**). While most plant ferredoxins had glutamate and aspartate residues that could interact with R101 and R140 on the Fe protein, they lacked positively charged residues on their surface and tended to form fewer salt-bridge interactions with E112 and E69 of the Fe protein. In general, our docking simulations suggest as the number of salt bridge interactions increased, the cofactor distance between a ferredoxin and the Fe protein decreased (**Fig. 5**). Our findings also indicate that nitrogen-fixing electron carriers form more complementary interactions with the Fe protein through salt-bridge interactions.

### Ferredoxins with cofactor distances ≤ 10 Å from the Fe protein can support nitrogenase activity in R. palustris

To validate that docking simulations could predict functional compatibility with nitrogenase, we replaced a native ferredoxin required for nitrogen fixation in *R. palustris* with heterologous ferredoxins that varied in their predicted cofactor distances from the Fe protein. To do this, we developed *R. palustris* as an expression platform for testing the effectiveness of heterologous ferredoxins *in vivo*. First, to ensure reliance on the heterologous ferredoxin, we deleted several other known electron carriers that might participate in delivering electrons to nitrogenase in *R. palustris*: *fldA*, the gene encoding flavodoxin that participates in electron transfer during iron limitation [40], and two 2[4Fe-4S] ferredoxins, *ferN* and *badB*. Next, we replaced *fer1* the primary 2[4Fe-4S] ferredoxin involved in nitrogen fixation in *R. palustris* [40], with genes encoding other 2[4Fe-4S] ferredoxins from a variety of bacterial species (**Table 1**). Ferredoxins from *Clostridium pasteurianum* (*Cp*), *Thermotoga maritima* (*Tm*)*, E. coli (EcFd)* and *Chlorobaculum tepidum* (*Ct*) were chosen because they have similar redox potentials to electron carriers involved in nitrogen fixation (Table 1). These ferredoxins were modeled with the Fe protein to measure the distance between the FeS clusters (**Table 1**). *Cp*Fd and *Tm*1175 were predicted to have the shortest electron transfer distances (10 Å or less) from the Fe protein, and the remaining bacterial ferredoxins have distances over 11 Å (**Table 1**). Only *Cp*Fd and *Tm*1175 were able to support growth under nitrogen-fixing conditions, consistent with their predicted shorter electron transfer distance (**Fig. 6**). Together, our results indicate that predicted cofactor distance can serve as a useful proxy for identifying ferredoxins compatible with nitrogenase and may guide the selection of electron carriers for nitrogen fixation in synthetic systems.

**Fig. 6.**
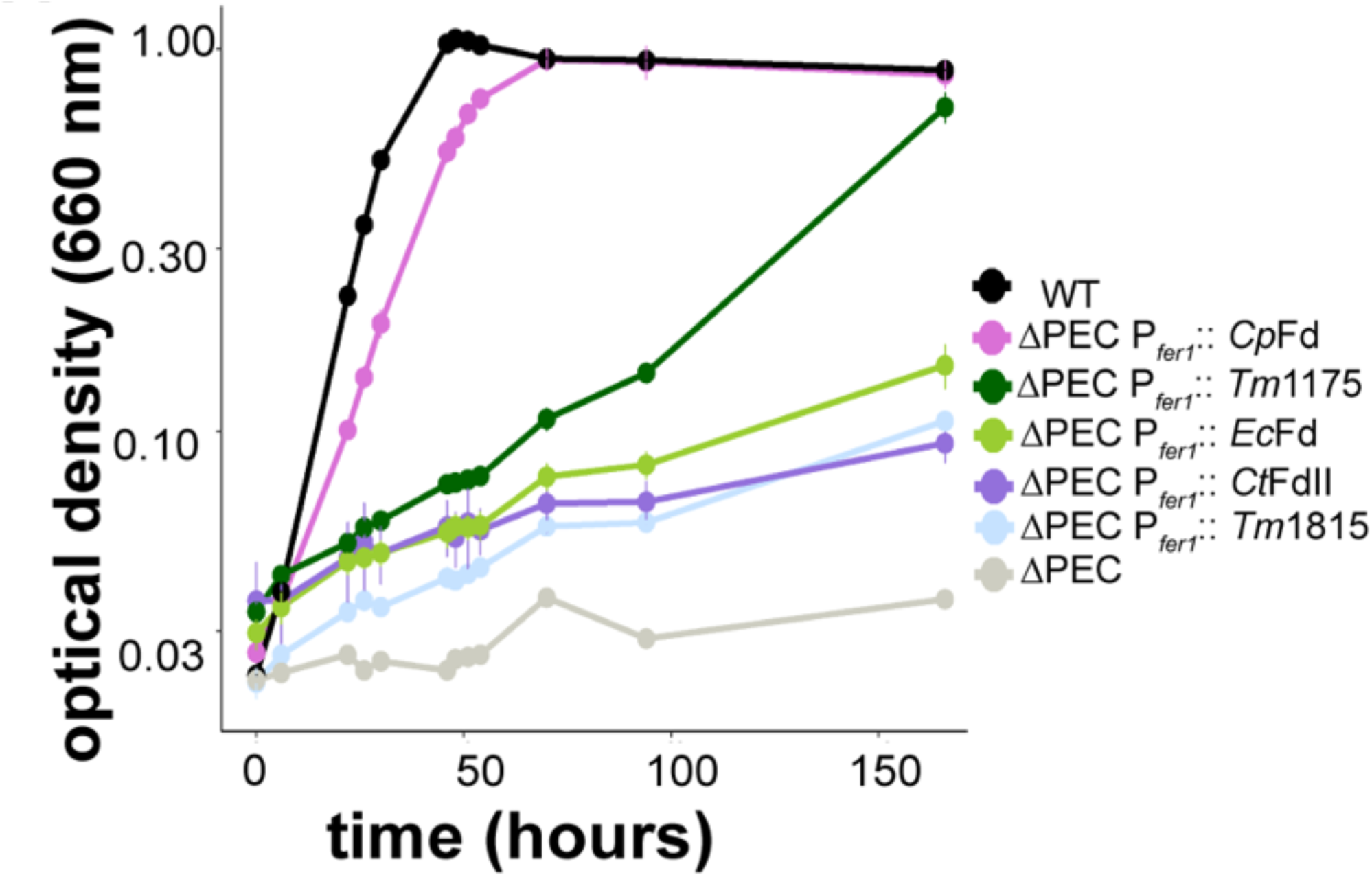
Heterologous expression of ferredoxins from *C. pasteurianum* or *T. maritima* support nitrogen fixation in an *R. palustris strain* lacking native protein electron carriers. Growth of *R. palustris* CGA009 (WT), *R. palustris* Δ*fer1* Δ*fldA* Δ*ferN* Δ*badB* (ΔPEC); or ΔPEC expressing a heterologous 2[4Fe-4S] ferredoxin from the *fer1* locus under nitrogen-fixing conditions, in the light, with 20 mM acetate. The ferredoxins include 2[4Fe-4S] ferredoxins from *C. pasteurianum* (*Cp*Fd), *T. maritima* (*Tm*1175 and *Tm*1815)*, C. tepidum* (*Ct*FdII), and *E. coli* (*Ec*Fd). Growth phenotypes are shown as the average of three biological replicates, and error bars represent one standard deviation from the mean.

**Table 1.** Properties of 2[4Fe-4S] ferredoxins heterologously expressed in *R. palustris* from the *fer1* locus.

| Organism and ferredoxin name | Average cofactor distance (Å) | Redox potential (mV)<sup>a</sup> | Electron tunneling rate, sec<sup>-1</sup> | Involved in nitrogen fixation |
| --- | --- | --- | --- | --- |
| *Clostridium pasteurianum*; CpFd | 8.9 | -380 | 4.58 x10<sup>7</sup> | Yes[69] |
| *Thermotoga maritima*; Tm1175 | 10.03 | -395, -490 | 8.63 x10<sup>6b</sup> | No |
| *Chlorobaculum tepidum*; CtFdII | 11.0 | -584 | 5.78 x10<sup>7</sup> | Yes[70] |
| *Escherichia coli*; EcFd | 11.4 | -418, -675 | 4.91 x10<sup>8b</sup> | No |
| *Thermotoga maritima*; Tm1815 | 26.1 | -320, -725 | ND | No |

Information on protein sequence and PDB ID for template used for these proteins are in supplemental table 5.
<sup>a</sup>A single potential indicates two clusters with the same potential.
<sup>b</sup> Higher redox potential was used to calculate electron tunneling rate for this ferredoxin.

## Discussion

The purpose of this study was to predict and potentially identify ways to improve the compatibility of a protein electron carrier with the Fe protein of nitrogenase. To achieve this, we simulated interactions between the Fe protein of *R. palustris* and a variety of electron carriers and validated our findings *in vivo*. Through our modeling, we found that ferredoxins and flavodoxins that play a role in nitrogen fixation had shorter distances between their electron-carrying cofactor and the [4Fe-4S] cluster in the Fe protein, falling within 10 Å (**Fig. 2**). This shorter distance likely reflects the ability of these electron carriers to associate with the Fe protein and enable fast electron tunneling (≥ 10^6^ sec^-1^), even when the redox potential of the electron carrier is higher than the Fe protein (**Fig. 3A** and **Supplemental Table 1**). In comparison, the distance between the [2Fe-2S] cluster of plant ferredoxins and the [4Fe-4S] cluster of the Fe protein exceeded 14 Å, the upper limit for efficient electron tunneling under physiological conditions (**Fig. 2**) [26, 28, 29, 50]. Even plant ferredoxins such as *At*Fd1, *At*Fd2, and *Zm*Fd1 that have redox potentials lower than Fe protein, have slower electron tunneling rates (≤ 10^6^ sec^-1^), because the cofactor distance is larger than 10 Å (**Supplemental Table 3**). Plant ferredoxins were shown for the first time to support some nitrogenase activity in an engineered *E. coli* when expressed them from an inducible promoter on a multi-copy plasmid [25]. However, the nitrogenase activity supported by these ferredoxins did not correlate with their redox potentials. Our findings may explain why these pant ferredoxins supported less nitrogenase activity, and this could be an important area of focus in future work.

We found that shorter distances between the electron-carrying cofactors tended to be inversely proportional to the number of electrostatic interactions between each docking complex (**Fig. 5**). Electrostatic interactions can enforce binding specificity with the partner proteins and help to distinguish between low affinity and high affinity complexes [51]. Further, electrostatic interactions have been implicated in determining specificity between ferredoxins and their binding partners, and there is experimental evidence to suggest that electrostatic interactions are specifically important for the ferredoxin-Fe protein interaction [42, 52–54]. Although protein modeling did not show that enhancing electrostatic interactions between ferredoxins and the Fe protein reduces their cofactor distance (data not shown), this may reflect a limitation of *in silico* protein design. We believe our observations in conjunction with previous reports could provide a framework for testing engineered variants of plant ferredoxins *in vitro* using purified nitrogenase or *in vivo* using chassis organisms like *R. palustris* or *E. coli* engineered to fix nitrogen [24, 25].

We also validated that our distance measurements reliably identified electron carriers that interact directly with nitrogenase (**Supplemental Table 1 and Fig. 2**). In *R. capsulatus*, *Rc*NifF, *Rc*FdB, *Rc*FdC, *Rc*FdD and *Rc*FdN are all upregulated under nitrogen-fixing conditions, but only *Rc*NifF and *Rc*FdN are known to donate electrons to nitrogenase while the other ferredoxins are implicated in other functions [41, 43, 55–58]. Consistent with these *in vivo* data, *Rc*NifF and *Rc*FdN exhibited the shortest modeled distances between their cofactor and the [4Fe-4S] cluster in the Fe protein. A similar pattern was observed for electron carriers from *R. palustris*, where only the electron carriers *Rp*NifF, *Rp*FdxB, *Rp*FerN, and *Rp*Fer1 are upregulated under nitrogen-fixing conditions [59]. Among these electron carriers, only *Rp*NifF, *Rp*FerN, and *Rp*Fer1 have been implicated in direct electron transfer to nitrogenase [40]. Docking models with *Rp*NifF, *Rp*FerN, and *Rp*Fer1 and the Fe protein all have cofactor distances 10 Å or less while docking models with *Rp*FdxB and the Fe protein have an average distance of 22.03 Å, well outside the range for efficient electron tunneling.

Moreover, this approach also suggested distinct roles for less-characterized electron carriers. For example, although both *Rc*FdN and *Rc*FdC are required for growth under nitrogen-fixing conditions, *Rc*FdC likely does not act as an electron donor for nitrogenase because of its high redox potential (around -285 mV) [41, 43]. The greater distance between the [2Fe-2S] cluster in *Rc*FdC and the [4Fe-4S] cluster in the Fe protein supports an alternative role for *Rc*FdC, perhaps as the electron donor for FprA, the nitrogen fixation-related flavoprotein, as has been previously proposed (**Fig. 2 and Supplemental Table 2**) [43]. Interestingly, *Rc*FdA and its homolog *Rp*FdxC have distances less than 10 Å in docking models with the Fe protein, and *Rc*FdA can act as electron donor to nitrogenase *in vitro* [60]. However, neither protein has been linked to nitrogen fixation *in vivo*, likely because they are not upregulated under these conditions and have roles outside of nitrogen fixation [45, 60]. These findings demonstrate that protein modeling can be a powerful tool for predicting the compatibility between electron carriers and the Fe protein, even when physiological function may be constrained by factors such as gene regulation. This approach may be particularly useful in diazotrophs, like *Methanosarcina acetivorans* and *Methanococcus maripaludis*, where it is still unclear what electron carriers are involved in electron transfer to nitrogenase.

To validate our observations from protein modeling that ferredoxins with cofactor ≤ 10 Å could support nitrogenase activity in a heterologous system, we tested this in *R. palustris*. *C. pasteurianum* is a known diazotroph, and *Cp*Fd was the first ferredoxin shown to deliver electrons to nitrogenase, and our result confirms that this ferredoxin can support nitrogenase activity [61]. In contrast, *T. maritima* lacks the genes for nitrogen fixation, and the physiological role of *Tm*1175 is unknown. However, *Tm*1175 is able to support *in vitro* activity of *T. maritima* MiaB, a radical S-adenosylmethionine methylthiotransferase [62]. The other ferredoxin we tested from *T. maritima*, *Tm*1815 had the largest edge to edge cofactor distance (26.1 Å) even though it is the most similar (28% amino acid identity) to a ferredoxin commonly found in *nif* gene clusters, FdxB, which also had a large, predicted cofactor distance, > 20 Å. While we have not ruled out other factors such as protein stability or lack of a specific ferredoxin-reducing enzyme affecting the activity of the 2[4Fe-4S] ferredoxins we tested, our data strongly support our observations from *in silico* modeling that distances between the cofactors could be used to identify heterologous ferredoxins that are compatible with nitrogenase.

To incorporate and maximize nitrogen fixation activity in non-nitrogen-fixing organisms will require understanding of how electrons will be delivered to nitrogenase. We have shown that *in silico* protein modeling offers a tool for identifying potential electron carriers that can supply this reducing power. Although redox potential is an important property of ferredoxins, it is not the sole determinant for compatibility between a ferredoxin and nitrogenase, and the distance between cofactors is likely a better predictor of which ferredoxin can support nitrogenase activity. Identifying ferredoxins or creating variants of existing ferredoxins capable of supporting nitrogenase activity could pave the way for introducing this pathway into organisms like plants that do not naturally fix nitrogen.

## Materials and method

### Protein models of ADP-bound Fe protein and electron carrier proteins

A structural model of ADP-bound Fe protein was built using the protein sequence from *R. palustris* (**Supplemental Table 4**). Initially, AlphaFold 2.0 was used to predict the Fe protein structure, but AlphaFold 2.0 generated only the ATP-bound conformation for the Fe protein homodimer. We used RoseTTAFold to generate a single monomer of the Fe protein and used the ADP-bound Fe protein structure from *A. vinelandii* (PDB entry 1FP6) as a template to generate the ADP-bound homodimer and incorporate the [4Fe-4S] cluster in PyMOL (v.3.0.3) [63],[64–66]. The structure of each bacterial and eukaryotic ferredoxin was generated using AlphaFold 2.0 in UCSF ChimeraX, and the PDB structures used to situate FeS clusters or flavin cofactors are shown in **Supplemental Table 1-3** [67]. The electrostatic potential of the surface of each ferredoxin was modeled using the Adaptive Poisson-Boltzmann Solver (APBS) computational tool in PyMOL [68] [64–66].

### Docking models of bacterial ferredoxins/flavodoxins and plant ferredoxins with NifH

*In silico* protein-protein docking models were prepared using the computational docking program ClusPro 2.0 using their default settings [37]. All docking simulations were prepared using the model of the *R. palustris* ADP-bound Fe protein described above as the receptor and the ferredoxin or flavodoxin as a ligand. The edge-to-edge distance between the Fe atom of cofactor in the ferredoxin or FMN of flavodoxin and the Fe atom of the [4Fe-4S] cluster of the Fe protein were measured in PyMOL using top three docking models that ClusPro 2.0 generated. The average edge-to-edge cofactor distance between cofactors, *R* (Å), of partner proteins using these models is represented in **Fig. 2** and **Supplemental Table 1 to 3**.

### Electron tunneling rate calculations

Electron tunneling rate refers to the frequency or probability at which an electron can transfer through a potential barrier. Factors that affect electron tunneling include driving force (ΔG^0^, mV) which is the difference between redox potentials of a ferredoxin and the Fe protein, and the edge-to-edge cofactor distance, *R* (Å) [29]. To calculate the electron tunneling rate (sec^-1^) from a ferredoxin or flavodoxin to ADP-bound Fe protein, ΔG^0^ was calculated using the redox potentials previously determined (see references in **Supplemental Tables 1-3**) and the edge-to-edge cofactor distance that we measured using protein-protein docking models.

The electron tunneling rate (sec^-1^) when ΔG^0^ was exergonic was calculated using **Eq. 1**:

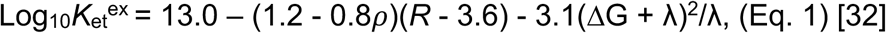

The electron tunneling rate (sec^-1^) when ΔG^0^ was endergonic was calculated using **Eq. 2**:

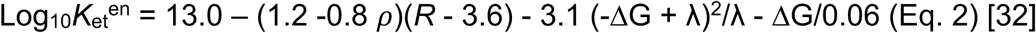

To calculate electron tunneling rate (sec^-1^), the reorganization energy (λ) used was 0.65 eV, and packing density (*ρ*) proposed to account for protein heterogeneity is 0.76 [29]. Reorganization energy (λ) is the energy required to disrupt the equilibrium of nuclear geometry of the partner proteins. The reorganization energies for tunneling electrons within proteins ranges between 0.6 eV to 0.9 eV in an aqueous phase, with an average value of 0.65 eV proposed for the reorganization energy within a cellular environment [29]. The calculated electron tunneling rates for each electron carrier is shown in **Fig. 3A** and **Supplemental Table 1-3**.

## Acknowledgements

This study was supported by the National Science Foundation CAREER award MCB 2338085. Support for K.T. was provided by award DE-SC0020252 from the U.S. Department of Energy, Office of Science, Basic Energy Sciences, Physical Biosciences program to K.R.F.

## Supplementary section contains the following information

**Supplemental materials and method**s: Culture conditions and genetic manipulation of *R. palustris* that were used in the study.

**Supplemental Table 1:** Information and template used for bacterial ferredoxins and flavodoxins involved in N-fixation including all data generated through the protein modeling.

**Supplemental Table 2:** Information, accession number, and template used for bacterial ferredoxins do not involve in N-fixation including all data generated through the protein modeling.

**Supplemental Table 3:** Information, accession number, and template used for plant ferredoxins do not involve in N-fixation including all data generated through the protein modeling.

**Supplemental Table 4:** Information and accession number of each Fe protein used for multiple sequence alignment in this study.

**Supplemental Table 5:** Information, accession number, and template used for 2[4Fe-4S] ferredoxins heterologously expressed from *fer1* promoter of *R. palustris*.

**Supplemental Fig. 1:** Multiple amino acid sequence alignments of the Fe protein homologs from different Nif clades.

**Supplemental Table 6:** Summary of primers, plasmids, and strains used for the study.

