## Supplemental methods, Supplemental tables 1-6, and supplemental figure 1 for "Predicting compatibility between ferredoxins and the Fe protein of nitrogenase using *in silico* protein modeling"

**Supplemental material and methods**

*Growth assays of R. palustris and its mutants*

Strains, plasmids, and primers are listed in **Supplemental Table 6**. All cultures of *R. palustris* were grown under anoxic conditions in sealed culture tubes with a headspace containing 97.5% N_2_ and 2.5% H_2_, with oxygen levels kept below 10 ppm. The cultures were maintained under photoheterotrophic conditions, using a 60 W incandescent bulb placed 5.5 inches from the tubes, providing a light intensity of 30 µmol photons m^-^² s⁻¹, as previously described in [1]. *R. palustris* strains were initially grown in minimal mineral medium [2], supplemented with 20 mM acetate and 0.1% yeast extract as carbon sources. Once cultures reached an optical density at 660 nm (OD_660_) of 1.1, they were diluted 1:100 into nitrogen-fixing medium, which is minimal mineral medium with no ammonium sulfate added and N_2_ as the only nitrogen source, with 20 mM acetate. The cells were allowed to undergo several doublings to acclimate the cells to nitrogen-fixing conditions and dilute any ammonium carried over from transfer. After the acclimation, cultures were used to inoculate fresh nitrogen-fixing medium. Growth was monitored over time by measuring optical density at 660 nm.

*Genetic manipulation of R. palustris*

In-frame deletion constructs of *fer1* (*rpa4631*), *fldA* (*rpa2117*), *ferN* (*rpa4629*), and *badB* (*rpa0662*) were prepared by PCR amplified fragments that included 1 kb of upstream of the start codon and 1 kb downstream of the stop codon. Genomic DNA from *R. palustris* CGA009 served as the template for amplification using Phusion High-Fidelity DNA polymerase (New England Biolabs). The two 1-kb fragments for each gene were ligated into pJQ200SK vector using *E. coli* DH5⍺-mediated assembly [3]. All plasmids were introduced into *R. palustris* by conjugation with *E. coli* S17-1, and double-crossover recombination events for gene deletions and screened using a previously described method [4]. The presence of deletions was confirmed by PCR.

Bacterial ferredoxin genes from *C. pasteurianum* (*Cp*), *T. maritima* (*Tm*), *C. tepidum* (*Ct*), and *E. coli* (*Ec*) were codon optimized for *R. palustris* using JAVA Codon Adaptation Tool (<https://www.jcat.de>) [5]. Replacement of *fer1* (*rpa4631*) with the bacterial ferredoxin genes was carried out by constructing a plasmid containing 25bp upstream of the *fer1* start codon and downstream of *fer1* stop codon fused to either side of the bacterial ferredoxin gene and ligated into pJQ200SK vector using *E. coli* DH5⍺-mediated assembly technique [3]. All plasmids were introduced into *R. palustris* by conjugation with *E. coli* S17-1, and double-crossover recombination events for gene deletions or allelic exchange were selected and screened using a previously described method [4]. Integration of bacterial ferredoxin genes was confirmed by colony PCR followed by Sanger sequencing. Strains were also validated using whole genome sequencing (SeqCenter).

**Supplemental Table 1. Nitrogen-fixing bacterial electron carriers used in this study.**

| **Bacterial ferredoxins/ flavodoxins** | **NCBI or UniProt accession number** | **PDB ID for template ^b^** | **Average cofactor distance (Å)^c^** | **Redox potentials (mV)** | **Calculated electron tunneling rate (sec^-1^) ^d^** |
| --- | --- | --- | --- | --- | --- |
| *Rp*Fer1 | P00207 | 2FGO [6] | 9.2 | -452,-583 [7] | 8.5 x 10^8^ |
| *Rp*FldA | Q6N7Y7 | 8V2Y [8] | 6.5 | -450 [8] | 3.8 x 10^9^ |
| *Rp*FerN | Q6N0Y0 | 1RGV [9] | 10.2 | ND^e^ | ND |
| *As*FdxH | P11053 | 1FRD [10] | 10.0 | -351 [11] | 5.4 x 10^6^ |
| *Ko*NifH | WP_004138775.1 | 1YOB [12] | 5.0 | -412 [13] | 2.7 x 10^10^ |
| *Rc*FdN | D5ARY6 | 1CLF [14] | 8.3 | -490 [15] | 1.87 x10^9^ |
| *Rc*FdA | D5AP15 | 6FD1 [16] | 6.9 | -419 [17] | 3.28 x10^9^ |
| *Rc*NifF | P52967 | 2WC1 [18] | 12.5 | -487 | 5.45 x10^8^ |

^a^ *Rhodopseudomonas palustris* (*Rp*), *Anabaena (Nostoc) sp.* PCC 7120 (*As*), *Klebsiella oxytoca* (*Ko*), *Rhodobacter capsulatus* (*Rc*).

^b^PDB template was selected based on the structural homology of proteins suggest by Phyre2[19]

^c^the edge-to-edge distance between the electron-carrying cofactor in the ferredoxin or flavodoxin and the [4Fe-4S] cluster in the Fe protein.

^d^ calculated from Eq. 1 and Eq. 2

^e^ND, not determined

**Supplemental Table 2. Non-nitrogen-fixing bacterial ferredoxins used in this study.**

| **Bacterial ferredoxins^a^** | **NCBI or UniProt accession number** | **PDB ID for template ^b^** | **Average cofactor distance (Å)^c^** | **Redox potentials (mV)** | **Calculated electron tunneling rate (sec^-1^) ^d^** |
| --- | --- | --- | --- | --- | --- |
| *Rp*BadB | Q6NC12 | 1RGV [9] | 12.6 | ND^e^ | ND |
| *Rp*FdxA | Q6N917 | 4ID8 [20] | 31.5 | ND | ND |
| *Rp*FdxB | Q6N0Z7 | 7QV7 [21] | 22.0 | ND | ND |
| *Rp*FdxC | Q6NCI3 | 1BQC [22] | 9.5 | ND | ND |
| *As*PetF | P0A3C8 | 1CZP [23] | 13.7 | -384 [11] | 8.9 x 10^5^ |
| *Cr*PetF | XP_001692808.1 | 1AWD [24] | 13.8 | -321 | 5.5 x 10^5^ |
| *Rc*FdC | D5ARY7 | 1FRR [26] | 14.9 | -285±10 | 9.81 x10^2^ |
| *Rc*FdB | D5ARX7 | 7QV7 [21] | 33.7 | ND | ND |
| *Rc*FdD | D5ANI4 | 8RHO [27] | 22.3 | ND | ND |
| *Rc*FdE | P80306 | 1E9M [28] | 14.2 | ND | ND |

^a^ *Rhodopseudomonas palustris* (*Rp*), *Anabaena (Nostoc) sp.* PCC 7120 (*As*), *Klebsiella oxytoca* (*Ko*), *Rhodobacter capsulatus* (*Rc*), *Chlamydomonas reinhardtii* (*Cr*).

^b^PDB template was selected based on the structural homology of proteins suggest by Phyre2 [19]

^c^ the edge-to-edge distance between the electron-carrying cofactor in the ferredoxin or flavodoxin and the [4Fe-4S] cluster in the Fe protein.

^d^ calculated from Eq. 1 and Eq. 2

^e^ND, not determined

**Supplemental Table 3. Plant ferredoxins used in this study.**

| **Plant ferredoxins^a^** | **NCBI or UniProt accession number** | **PDB ID for template ^b^** | **Average edge to edge cofactor distance (Å)^c^** | **Redox potentials (mV)** | **Calculated electron tunneling rate (sec-^1^) ^d^** |
| --- | --- | --- | --- | --- | --- |
| *Zm*FDI | P27787.1 | 5H57 [29] | 16.6 | -423 [6] | 1.3 x 10^4^ |
| *Zm*FDII | O80429.1 |  | 15.6 | -406 [30] | 2.2 x 10^4^ |
| *Zm*FDIII | NP_001105346.1 |  | 14.7 | -321 [31] | 1.2 x 10^4^ |
| *At*FD1 | NP_172565.1 | 4ZHO [32] | 16.1 | -425 [33] | 1.6 x 10^4^ |
| *At*FD2 | NP_176291.1 |  | 15.8 | -433 [33] | 2.6 x 10^4^ |
| *At*FD3 | NP_180320.1 |  | 13.7 | -337 [33] | 4.7 x 10^5^ |
| *At*MFD1 | NP_001329852 | 5UJ5 [34] | 15.2 | ND^e^ | ND |
| *At*MFD2 | NP_001031685 | 2WLB [35] | 15.9 | ND | ND |
| *Os*FD1 | NP_001390240.1 | 1GAQ [36] | 15.3 | ND | ND |
| *Os*FD4 | XP_066166407.1 | 1CZP [23] | 13.6 | ND | ND |
| *Ta*FD4 | NP_001413721.1 | 1GAQ [36] | 10.7 | ND | ND |

^a^ *Arabidopsis thaliana* (*At*), *Zea mays* (*Zm*), *Triticum aestivum* (*Ta, Ta*Fd4), *Oryza sativa* (*Os*).

^b^PDB template was selected based on the structural homology of proteins suggest by Phyre2[19]

^c^ the edge-to-edge distance between the electron-carrying cofactor in the ferredoxin or flavodoxin and the [4Fe-4S] cluster in the Fe protein.

^d^ calculated from Eq. 1 and Eq. 2

^e^ND, not determined

**Supplemental Table 4. NifH sequences used for multiple sequence alignment in this study**

| **Organism** | **Fe protein** | **NCBI or UniProt accession number** |
| --- | --- | --- |
| *Rhodopseudomonas palustris* | NifH | WP_011160152 |
| *Azotobacter vinelandii* | NifH | WP_012698831.1 |
| *Anabaena PCC 7120* | NifH | AAB65801.1 |
| *Klebsiella pneumoniae* | NifH | P00458 |
| *Rhodospirillum rubrum* | NifH | WP_011388765.1 |
| *Rhodobacter capsulatus* | NifH | WP_013066316.1 |
| *Chlorobaculum tepidum* | NifH | WP_010933198.1 |
| *Clostridium pasteurianum* | NifH | WP_003447877.1 |
| *Methanosarcina acetivorans* | NifH | WP_011023791.1 |
| *Rhodopseudomonas palustris* | AnfH | WP_011157001.1 |
| *Rhodobacter capsulatus* | AnfH | WP_013066329.1 |
| *Azotobacter vinelandii* | AnfH | P16269.1 |
| *Rhodopseudomonas palustris* | VnfH | WP_011156939.1 |
| *Azotobacter vinelandii* | VnfH | P15335.1 |
| *Methanococcus maripaludis* | NifH | WP_011170797.1 |

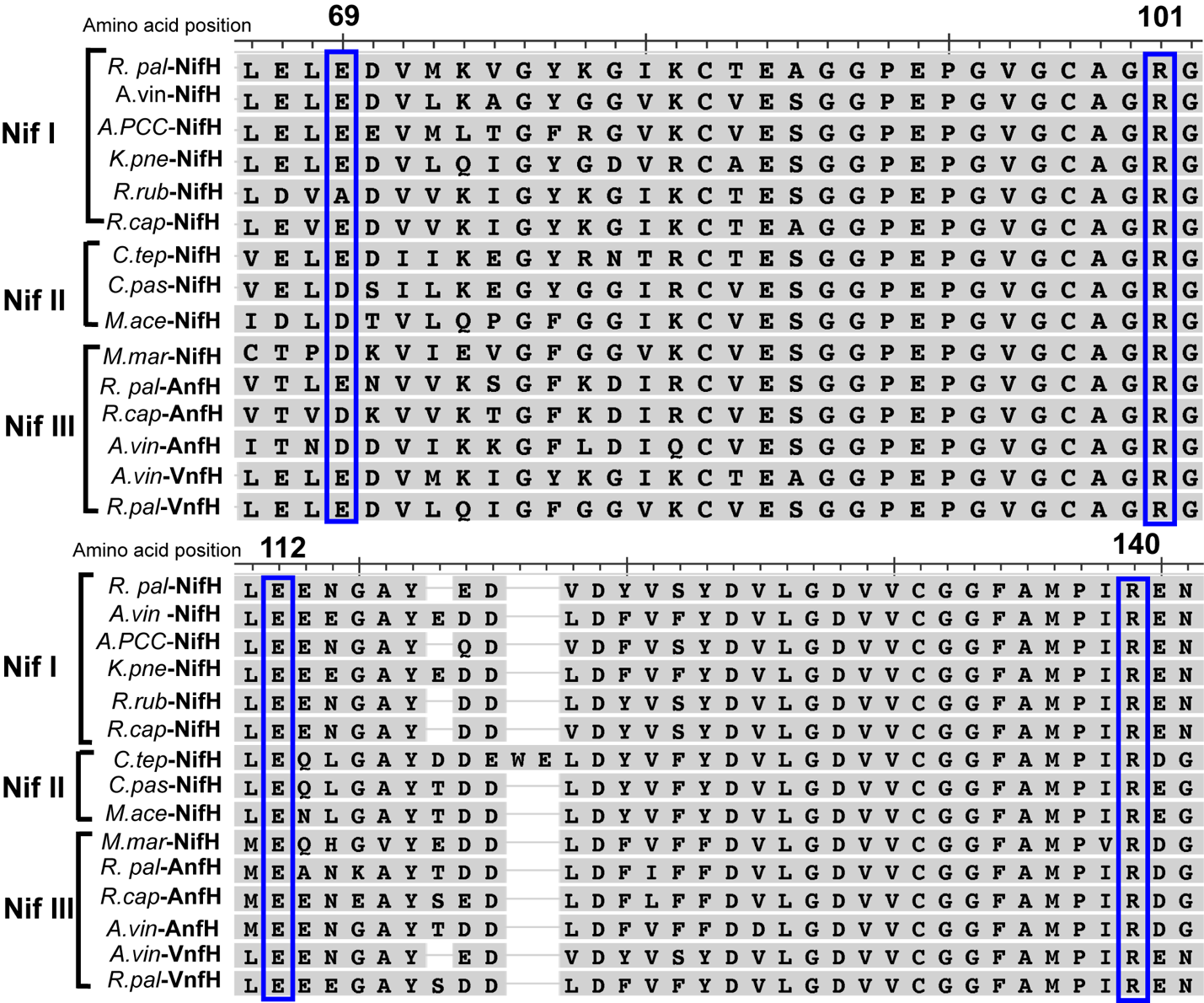

**Supplemental Fig. 1. Multiple amino acid sequence alignments of Fe protein homologs from Nif clades.** Charged residues including arginine (R) 101, glutamine (E) 112, and arginine (R) 140 are 100% conserved; residue glutamine (E) 69 is 70% conserved or replaced with mostly aspartate (D) among Fe proteins of Nif clades (Nif I, Nif II, and Nif III) [37]. COBALT (NCBI) [38] was used to align amino acid sequences of Nif, Vnf, or Anf homologs from selected model organisms including *R.pal*: *Rhodopseudomonas palustris*, *A.vin*: *Azotobacter vinelandii*, *K.pne*: *Klebsiella pneumoniae*, *R.rub*: *Rhodospirillum rubrum*, *R.cap*: *Rhodobactor capsulatus*, *C.tep*: *Chlorobaculum tepidum*, *C.pas*: *Clostridium pasteurianum*, *M.ace*: *Methanosarcina acetivorans, M.mar*: *Methanococcus maripaludis*.

**Supplemental table 5: 2[4Fe-4S] ferredoxins heterologously expressed in *R. palustris*.**

| **Organism** | **Ferredoxin name** | **NCBI accession number** | **PDB ID for template** |
| --- | --- | --- | --- |
| *Clostridium pasteurianum* | *Cp*Fd | P00195.2 | 1CLF[14] |
| *Thermotoga maritima* | *Tm*1175 | WP_004080188.1 | 1RGV[9] |
| *Chlorobaculum tepidum* | *Ct*FdII | WP_010932930.1 | 1RGV[9] |
| *Esherichia coli* | *Ec*Fd | WP_001196283.1 | 2ZVS[39] |
| *Thermotoga maritima* | *Tm*1815 | WP_004082361.1 | 2ZVS[39] |

**Supplemental Table 6: Primer, plasmids, and strains used for this study.**

| **Primer name** | **Primer sequence (5’ to 3’)** | **Reference** |
| --- | --- | --- |
| pJQ200SK_vecF | CGCTGATCTCCGAAGCCGAGCACGGGCTCTAGAACTAGTGGATCCCCGG | [40] |
| pJQ200SK_vecR | TCGAATTCCTGCTCCTGCAAGACGCCTCCAGCTTTTGTTCCCTTTAGTGAGGG | [40] |
| Cp_usF | ATGGCCTACAAGATCGCCGACTCGTG | This study |
| Cp_dsR | TTACTCCTGGACCGGGGCGC | This study |
| CtFdII_usF | ATGGCCCTGTACATCACCGAGGAG | This study |
| CtFdII_dsR | TTAGCCCTGGACGATGCACTCGG | This study |
| Ec_usF | ATGGCCCTGCTGATCACCAAGAAG | This study |
| Ec_dsR | TTAGATCTTGTCGGCGTGGTGCATC | This study |
| Tm1175_usF | ATGGCCAAGAACTGGTACCCGGTC | This study |
| Tm1175_dsR | TTAGCCGTCGGCCGAGACCTC | This study |
| Tm1815_usF | ATGGCCGAGGCCAAGAACGCCCC | This study |
| Tm1815_dsR | TTACGGCTCCGGCTTGGTCTCGGTCTCC | This study |
| **Plasmid** | **Description, host strain, and selective marker** | **Reference** |
| p*fer1::*Cpas Fd | pJQ200Sk vector *C. pasteurianum* Fd with 25bp upstream and downstream of *fer1* in S17-1 *E. coli*, gentamycin 20µg/mL | This study |
| p*fer1::*Ctep Fd | pJQ200Sk vector *C. tepidum* Fd with 25bp upstream and downstream of *fer1* in S17-1 *E. coli*, Gentamycin 20µG/mL | This study |
| p*fer1::*Ec Fd | pJQ200Sk vector expressing *E. coli* Fd with 25bp upstream and downstream of *fer1* in S17-1 *E. coli*, Gentamycin 20µG/mL | This study |
| p*fer1::*TM1175 Fd | pJQ200Sk vector expressing *T. maritima* TM1175 Fd with 25bp upstream and downstream of *fer1* in S17-1 *E. coli*, Gentamycin 20µG/mL | This study |
| p*fer1::*TM1815 Fd | PJQ200Sk vector expressing *T. maritima* TM1815 Fd with 25bp upstream and downstream of *fer1* in S17-1 *E. coli*, Gentamycin 20µG/mL | This study |
| **Strains** | **Description** | **Reference** |
| WT | CGA009 | [41] |
| ∆PEC | Deletion of electron carrier protein encoding genes – *fer1*, *fldA*, *ferN*, and *badB* | This study |
| ∆PEC p*fer1*::*Ec* Fd | ∆PEC strain with *E. coli* Fd expressed at the *fer1* locus | This study |
| ∆PEC p*fer1*::Ctep Fd | ∆fer1ferNfldAbadB strain with *C. tepidum* Fd expressed at the *fer1* locus | This study |
| ∆PEC p*fer1*::Cpas Fd | ∆PEC strain with *C. pasteurianum* Fd expressed at the *fer1* locus | This study |
| ∆PEC p*fer1*::*Tm*1175 Fd | ∆PEC strain with *T. maritima* TM1175 Fd expressed at the *fer1* locus | This study |
| ∆PEC pfer1::*Tm*1815 | ∆PEC strain with *T. maritima* TM1815 Fd expressed at the *fer1* locus | This study |
